# Intrinsically disordered linkers control tethered kinases via effective concentration

**DOI:** 10.1101/2020.04.04.023713

**Authors:** Mateusz Dyla, Magnus Kjaergaard

## Abstract

Kinase specificity is crucial to the fidelity of signalling pathways, yet many pathways use the same kinases to achieve widely different effects. Specificity arises in part from the enzymatic domain, but also from the physical tethering of kinases to their substrates. Such tethering can occur via protein interaction domains in the kinase or via anchoring and scaffolding proteins, and can drastically increase the kinetics of phosphorylation. However, we do not know how such intra-complex reactions depend on the link between enzyme and substrate. Here we show that the kinetics of tethered kinases follow a Michaelis-Menten like dependence on effective concentration. We find that phosphorylation kinetics scale with the length of the intrinsically disordered linkers that join the enzyme and substrate, but that the scaling differs between substrates. Steady-state kinetics can only partially predict rates of tethered reactions as product release may obscure the rate of phospho-transfer. Our results suggest that changes in signalling complex architecture not only enhance the rates of phosphorylation reactions, but may also alter the relative substrate usage. This suggests a mechanism for how scaffolding proteins can allosterically modify the output from a signalling pathway.

## Introduction

Cellular signalling relies on cascades of enzymes such as protein kinases and phosphatases. Signalling fidelity requires kinases to use specific substrates, although the same kinases can take part in different pathways with different substrates. For example, protein kinase A (PKA) is involved in signalling pathways throughout cell biology.^1^ Kinase specificity arises in part from the linear motifs surrounding the phospho-accepting residue. However, *in vivo* substrate usage can only be predicted partially from sequences motifs as kinase signalling is also regulated through the local abundance of enzymes.^2^ The latter class of effects is much less understood quantitatively.

The local abundance of signalling enzymes is regulated at many levels. At the cellular level, kinases are targeted to e.g. an organelle or the cell membrane. At the molecular level, enzymes are targeted to a subset of their potential substrates by tethering to scaffolding proteins. In multidomain enzymes, such tethering occur via protein interaction domains attached to the catalytic domain.^3^ Alternatively, enzymes can be tethered by scaffolding proteins that bind the enzyme and substrates non-covalently.^4^ PKA is regulated by more than 50 different A kinase anchoring proteins (AKAPs) that direct the kinase to different pathways.^5^ PKA may dissociate from its regulatory subunit and thus AKAPs upon activation, but recently it was demonstrated that active PKA may remain tethered to AKAP.^6^ The intrinsically disordered AKAPs thus restrict the kinase to a region whose dimensions are defined by the length and compactness of the anchoring protein.^7^ Similarly, disordered linkers of variable length regulate the activity of the dodecameric kinase CaMKII by controlling the balance between inter- and intra-subunit phosphorylation.^8^ Many kinase signalling reactions thus occur internally in proteins or in protein complexes.

When a kinase is physically associated to its substrate, the tethered substrate is distinct from all other substrates in the reaction mixture and is usually phosphorylated much faster. The kinetics of such reactions differ from steady-state enzyme kinetics as tethered catalysis is effectively single turnover. Due to the switch-like functions of many kinases, they may only need to phosphorylate a single substrate to exert a biological function.^9^ This can be seen e.g. in the negative feedback loop formed by AKAP79-tethered PKA, which inactivates adenylate cyclase by phosphorylation.^10^ The rate of intramolecular (or intra-complex) reactions is expected to be concentration independent although this is rarely tested for tethered kinases. Instead, the encounter rate is determined by the connection between enzyme and substrate. The quantitative description of tethered catalysis is hindered by the lack of framework to describe the connection in kinetic terms. We propose that many linkers can be accounted for by the effective concentration of substrate they enforce. Enzyme scaffolding proteins and interdomain linkers in kinases are often intrinsically disordered as they allow the kinase domain to search for its substrates.^11^ For fully disordered linkers, the effective concentrations can be estimated from power law scaling with the length of the linker.^12^ Kinases tethered by disordered protein linkers thus offer a chance to understand the general principles that govern tethered catalysis.

Here we investigate how the connection between a kinase and its substrate affects the phosphorylation kinetics. We use a model system consisting of the catalytic domain of PKA tethered to a substrate via variable intrinsically disordered linkers. We show that phosphorylation rate varies by several orders of magnitude depending on the substrate and the linker. We show that phosphorylation rate follows a Michaelis-Menten like dependence on the effective concentration enforced by the linker, with steady-state parameters partially predicting the rates of tethered single turnover phosphorylation. This provides a baseline for interpreting how signalling complex architecture regulates tethered enzymes.

## Materials and Methods

### Preparation of DNA constructs

The plasmid containing the catalytic domain of PKA (PKAc) was a gift from Susan Taylor via Addgene (#14921).^13^ The other plasmids were synthesized by Genscript and codon optimized for expression in *E. coli*. The coding regions were cloned into pET15b vectors using the NdeI/XhoI sites. The disordered GS linkers from the linker library generated previously^12^ were cloned into the protein constructs using unique NheI/KpnI sites. The sequences of protein constructs are given in the SI.

### Protein expression and purification

Protein constructs containing the catalytic domain of PKA linked by variable length linkers to the MBD2 coiled-coil domain (MBD2-(GS)_n_-PKAc) were expressed in C41(DE3) cells in LB medium with 100 µg/mL ampicillin at 37°C and shaking at 120 RPM. The cultures were induced with 1 mM IPTG at OD_600_=0.6-0.8 and the temperature was decreased to 30°C for overnight expression. Protein constructs containing PKA substrates linked to the p66α coiled-coil domain (p66α-(GS)_n_-substrate) were expressed in BL21(DE3) cells in ZYM-5052 auto-induction medium^14^ containing 100 µg/mL ampicillin at 37°C and shaking at 120 RPM overnight.

The cells were harvested by centrifugation (15 min, 6000 g), bacterial pellets were resuspended in binding buffer (20 mM NaH_2_PO_4_, 0.5 M NaCl, 5 mM imidazole, 0.1 mM TCEP, 0.2 mM PMSF, 50 mg/L of leupeptin, 50 mg/L pepstatin, 50 mg/L chymostatin, pH 7.4) and lysed by sonication (50% duty cycle, maximum power of 75%, sonication time of 6 min). The lysate was centrifuged (20 min, 27000 g) and the supernatant was applied to gravity flow columns packed with Ni-NTA Superflow (QIAGEN). The columns were washed with buffers containing increasing concentrations of imidazole (20-50 mM) and were eluted with elution buffer (20 mM NaH_2_PO_4_, 0.5 M NaCl, 500 mM imidazole, 0.1 mM TCEP). Eluted protein samples containing PKA substrates were dialyzed against binding buffer without imidazole overnight, followed by cleavage of His-tag by thrombin (1:1000 thrombin to substrate ratio) at 37°C and shaking at 300 RPM for 30 min. Cleaved samples were applied to gravity flow columns packed with Ni-NTA Superflow (QIAGEN) and flow-through was collected. All protein samples were additionally purified by size-exclusion chromatography (SEC) in TBS buffer (20 mM Tris-base, 150 mM NaCl, 0.1 mM TCEP, pH 7.6) using Superdex 200 Increase and Superdex 75 Increase columns (GE healthcare) to purify PKAc and its substrates, respectively.

### Quench-flow measurements

Single-turnover kinetic measurements were performed using a KinTek Corporation RQF-3 Rapid quench-flow instrument at 30°C. MBD2-(GS)_n_-PKAc and p66α-(GS)_n_-substrate were diluted to 10x final concentration in enzyme dilution buffer (50 mM Tris-base, 0.1 mM EGTA, 1 mg/ml BSA, 1 mM TCEP, pH 7.6) and TBS (20 mM Tris-base, 150 mM NaCl, pH 7.6), respectively. Quench flow experiments were executed by loading the catalytic domain of PKA and its substrates (final concentration 0.5 µM unless indicated otherwise) into one sample loop and [γ-^32^P]ATP (final concentration 0.1 mM of 100-200 counts per minute (c.p.m.) pmol^-1^) into the other. Phosphorylation reactions were carried out in the reaction buffer composed of 50 mM Tris-base, 0.1 mM EGTA, 10 mM magnesium acetate, 150 mM NaCl, pH 7.6. The reactions were quenched using 150 mM phosphoric acid using the “constant quench” option to limit the total quenched reaction volume to 70 µL. Phosphorylated substrates were separated from unreacted ATP by spotting the quenched reactions onto P81 phosphocellulose disks (Jon Oakhill, St. Vincent Institute, Melbourne). The filters were washed 3 times with 75 mM phosphoric acid, rinsed with acetone, dried, and counted on the ^32^P channel in scintillation counter as c.p.m. Control experiments were performed to determine background phosphorylation level of substrate motifs in the absence of PKAc, as well as autophosphorylation of PKAc in the absence of substrates. Background values (typically negligible) were subtracted from the raw data. Specific phosphorylation was determined as amount of ^32^P-incorporated substrate (pmol) based on ^32^P-ATP standard curve, where c.p.m. values of known amounts of spiked ^32^P-ATP were measured.

### Steady-state kinetics

PKAc (without the MBD2 dimerization domain) was diluted to 10x final concentration in enzyme dilution buffer (50 mM Tris-base, 0.1 mM EGTA, 1 mg/ml BSA, 1 mM TCEP, pH 7.6) and p66α-(GS)_n_-substrate was diluted to 5x final concentration in TBS (20 mM Tris-base, 150 mM NaCl, pH 7.6). Steady-state experiments were executed by manual addition of [γ-^32^P]ATP (final concentration 0.1 mM of 100-200 c.p.m. pmol^-1^) into reaction mix containing 1nM PKAc and p66α-(GS)_n_-substrate (final concentration 10-1820 µM) in a reaction buffer (50 mM Tris-base, 0.1 mM EGTA, 10 mM magnesium acetate, 150 mM NaCl, pH 7.6) at 30°C. Every minute 5 µl of the reaction mix was spotted onto a P81 filter disk and placed into 75 mM phosphoric acid to quench the reaction. The filters were washed 3 times with 75 mM phosphoric acid, rinsed with acetone, dried, and counted on the ^32^P channel in scintillation counter as c.p.m. Control experiments were performed in the same way as in quench-flow experiments, and specific phosphorylation was determined likewise. Initial velocities at different substrate concentrations were derived from a slope of a linear regression as pmol of ^32^P-incorporated substrate per minute. The data were then fitted by a Michaelis-Menten model, and *k*_cat_ was calculated by dividing *V*_max_ by 0.005 pmol of PKAc and by 60 (min/s).

## Results

To probe the linker dependence of tethered catalysis, we designed a model system composed of the catalytic domain of protein kinase A (PKAc) tethered to its substrate via disordered linkers (Fig. 1A, S1). We used the optimal PKA substrate Kemptide (WT in the following)^16^ and varied substrate quality by mutagenesis. The kinase and substrate were made as separate proteins and joined non-covalently using heterodimeric coiled-coil domains. We use the anti-parallel coiled-coil formed between p66α and MBD2^15^ as it has low nM affinity, and is stable on the timescale of the tethered reaction. The coiled-coil domains were joined to the PKAc/substrate by an exchangeable linker flanked by restriction sites compatible with our library of disordered linkers.^12^ To approximate passive, entropic spacers, we used glycine-serine repeat linkers containing 2, 20, 60 and 120 residues.

**Figure 1:**
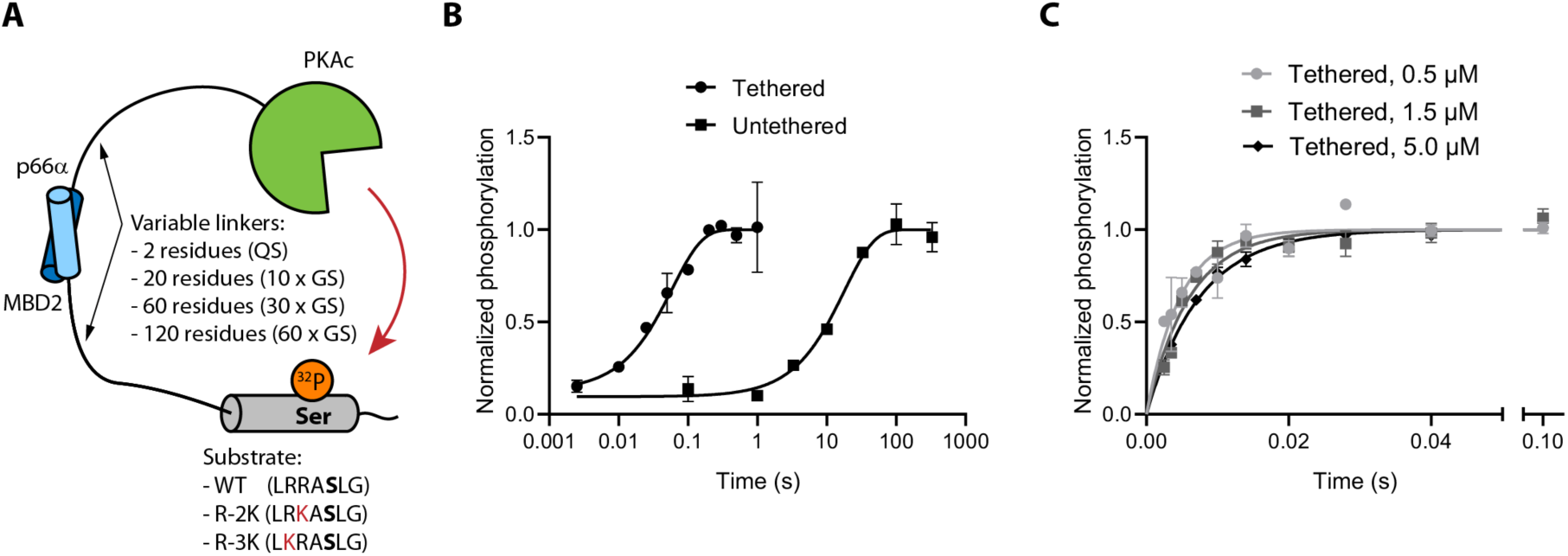
A reductionist model system for tethered phosphorylation. (A) The catalytic domain of PKA and its consensus substrate motif are joined to the coiled-coil domain from MBD2 and p66α, respectively. MBD2 and p66α form an anti-parallel heterodimer with nanomolar affinity.^15^ Substrate quality was varied by mutation away from the consensus motif, and the linker was varied by including GS-repeats of variable length. (B) Quench-flow was used to measure single-turnover kinetics by following incorporation of ^32^P into the substrate. Tethering of enzyme using combined linker length of 40 residues increased the rate by ∼300-fold at a substrate and enzyme concentrations of 0.5 µM. (C) The rate of the tethered catalysis decreased slightly with increasing concentration of substrate and enzyme.

We followed single-turnover phosphorylation in an equimolar mix of kinase and substrate using quench-flow. The reaction was started by adding [γ-^32^P]ATP, quenched by phosphoric acid at reaction times from 2.5 ms to 330 s, and followed from the amount of ^32^P in the substrate. Initially, we compared identical tethered and untethered reactions, where tethering increased reaction rate by more than 100-fold (Fig. 1B). Intra-complex reactions are concentration independent, whereas the rate of bimolecular reactions increases with concentration. Therefore, we tested how the concentration of the enzyme and substrate affected the phosphorylation rate. While remaining below the *K*_M_ of the substrate, the concentration was increased 10-fold, which led to a slightly slower phosphorylation (Fig. 1C). These observations suggest that phosphorylation occurs intra-complex with negligible contributions from trans-phosphorylation.

To test how linkers affect phosphorylation, we varied the lengths of the GS repeats. By combining kinase and substrate variants, we created 6 complexes with a combined linker length spanning from 20 to 180 disordered residues in addition to the 17 residues from restriction sites and the short extra linker in the substrate. For all variants the phosphorylation reaction could be described well with a single exponential model, where the phosphorylation rate decreased monotonically with increased linker length (Fig. 2A, S2). This suggests that kinase linkers and anchoring proteins directly regulate intra-complex kinase reactions.

**Figure 2:**
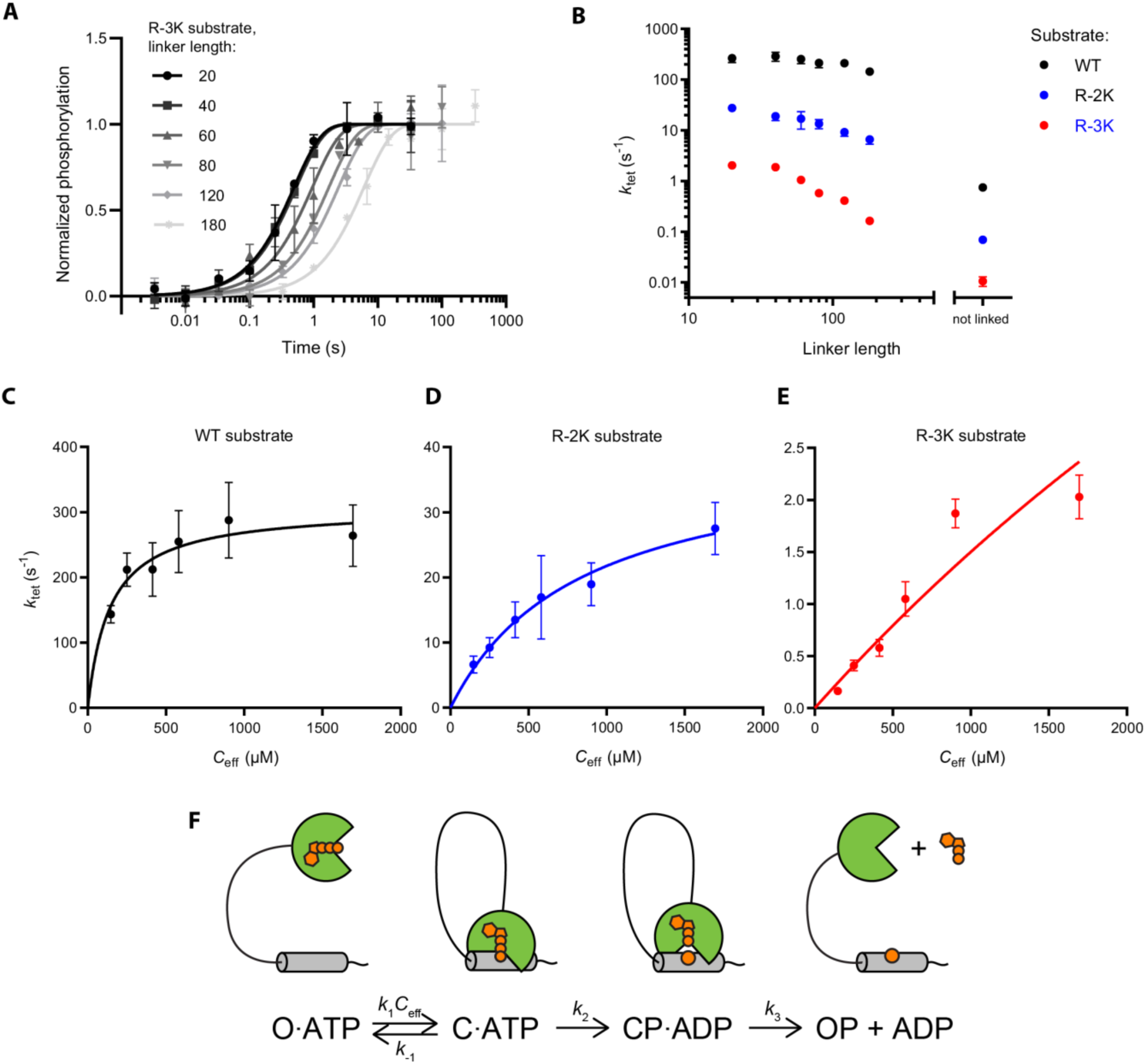
Linker dependence of single-turnover phosphorylation rates. (A) Quench-flow kinetics of the phosphorylation reactions of the R-3K substrate with different linker lengths. The linker length is the combined number of GS-repeats in the two linkers. (B) The observed rate of phosphorylation of three different substrates as a function of the combined linker length. (C-E) Phosphorylation rates as a function of the estimated effective concentration enforced by the linker. Effective concentrations are estimated from the power law measured previously for GS-linkers^12^ assuming that the presence of the coiled-coil domain is negligible. (F) Kinetic scheme of the tethered system under saturating concentration of ATP. O·ATP – open state of the tethered system, ATP-bound; C·ATP – closed catalytically competent state, ATP-bound; CP·ADP – closed phosphorylated state, ADP-bound; OP – open, nucleotide-free state. Nucleotide and phosphorylation are shown as orange moieties in the cartoon.

Next, we tested whether substrate quality affects the linker dependence of phosphorylation. PKA recognizes positive residues in positions −2 and −3 relative to the phosphorylation site with a preference for arginine over lysine.^16^ To weaken the site, we mutated each of the two arginine residues in the substrate motif to lysine resulting in variants R-2K and R-3K. The phosphorylation rate decreased monotonically with linker length for both mutants, although with different magnitudes (Fig. 2B). Phosphorylation of the WT substrate only slows 2-fold from the shortest to the longest linkers, whereas the R-3K substrate slows by more than 10-fold. This suggests that substrates are affected differently by tethering, and that the properties of the linker and substrate should be considered together.

The attenuated linker dependence of the WT substrate could in principle be caused by ATP binding becoming the limiting factor at high phosphorylation rates. To test this possibility, we carried out quench flow experiments at a 1 mM ATP instead of 100 µM. The amount of [γ-^32^P]ATP was kept constant to stay within permitted radioactivity levels, and so we increased the protein concentration to 5 µM, which had a minimal effect on kinetics (Fig. 1C), to maintain a robust signal. For the WT substrate, phosphorylation rates increased by ∼30% (Fig. S3) for all linker combinations. A uniform rate enhancement at 1 mM ATP is expected if the kinase is not fully saturated with ATP at 100 µM. Previous studies report a *K*_M_ for ATP of 10 µM at different buffer condition,^17^ so the kinase may not be fully saturated at 100 µM. To test this explanation, we also tested a slower R-3K variant with 1 mM ATP, and saw a similar increase in the observed rate (Fig. S2) consistent with an increase in level of bound ATP. This suggests that the attenuated linker dependence for the WT is not due to ATP binding becoming kinetically limiting, but rather represents saturation of the tethered substrate.

Tethered reactions are expected to follow first order kinetics with the contact frequency of reactants determined by their physical connection.^18^ The contact frequency can be expressed as an effective concentration, which in our case corresponds to the concentration of free substrate that would encounter the enzyme as often as the tethered substrate. Such effective concentration can occasionally be measured by competition experiments,^12,19,20^ although in most proteins has to be estimated theoretically.^21–23^ Our group previously measured the scaling of effective concentrations with the length of disordered linkers including the sequences used here.^12^ To estimate the effective concentration of the tethered substrate, we used the power law determined for GS-linkers previously, and plotted phosphorylation rates as a function of effective concentration (Fig. 2C-E). The tether in our kinase-substrate system also includes the coiled-coil domain, which act as a rigid link in the flexible linker. Previous effective concentration measurements suggest a similar scaling with linker length for such a system, so we disregard the contribution of the coiled-coil domain although it potentially introduces a systematic error.^19^

The phosphorylation rate saturated at high effective concentration for all variants. The WT substrate was close to saturation already at the concentrations enforced by the longest linkers. In contrast, R-2K only saturated at the highest effective concentration enforced by the shortest linkers, whereas R-3K has not plateaued yet at the highest effective concentration. The maximal rate also differed strongly between the substrates, as the R-2K and R-3K variants saturated at much lower rates than the WT substrate. In total, this suggests that the single turnover reaction can be described by a Michaelis-Menten like dependence on the effective concentration with a halfway saturation point and maximal turnover characteristic of each substrate, where *k*_tet,max_ is the maximum phosphorylation rate and *C*_eff,50_ is an effective concentration at half-saturation:

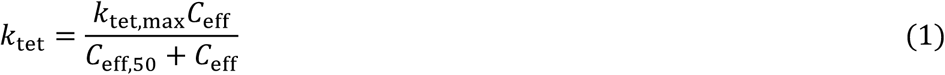

To describe the kinetics of tethered phosphorylation reactions, we derived rate equations (SI text) based on a product-release limiting model for PKA (Figure 2F).^17,24^ Under the assumption that phosphorylation (*k*_2_) and product release (*k*_3_) are irreversible and the closed, catalytically competent complex (C) is in a rapid equilibrium with an open form (O), the rate of phosphorylation in the tethered system (*k*_tet_) follows the following relationship on effective concentration (*C*_eff_):

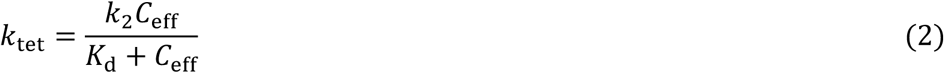

The rate of phosphotransfer (*k*_2_) is directly obtained from the maximum rate of phosphorylation in the tethered system (*k*_tet,max_) and *K*_d_ is obtained from an effective concentration at half-saturation *C*_eff,50_, based on the fit of experimental data to Equation 1. The fitting parameters are given in Table 1. *K*_d_ determined in this way for the WT substrate is equal to 142 µM, similar to the literature value of 200 µM^17^, and it increased to 837 µM and 8400 µM for R-2K and R-3K, respectively.

**Table 1:** Kinetic parameters obtained from fits of experimental data.

|  |  | WT | WT | R-2K | R-3K |
| --- | --- | --- | --- | --- | --- |
|  |  | 1mM ATP |  |  |  |
| **Tethered** | $k_{\text{tet,max}} (\text{s}^{-1})$ | 415 ± 47 | 307 ± 21 | 40.0 ± 3.5 | = 14.1 |
| | $C_{\text{eff},50} (\mu\text{M})$ | 107 ± 59 | 142 ± 42 | 837 ± 150 | 8400 ± 910 |
| **Untethered** | $k_{\text{cat}} (\text{s}^{-1})$ | | 37.3 ± 0.8 | 33.5 ± 0.4 | 14.1 ± 0.3 |
| | $K_M (\mu\text{M})$ | | 24.0 ± 2.1 | 200 ± 5.2 | 973 ± 43 |

To test whether the rates of tethered catalysis can be predicted from steady-state kinetics, we recorded a matching untethered steady-state enzyme kinetics for each substrate (Fig. 3). The *K*_M_ of the untethered reaction increased from 24.0 µM in the WT substrate to 200 and 973 µM in R-2K and R-3K, respectively. *K*_M_ values for all substrates are an order of magnitude below the corresponding *K*_d_ values. To quantify the relationship between *K*_M_ and *K*_d_, we derived steady-state parameters as a function of rate constants in untethered system (SI text), which resulted in the following equation:

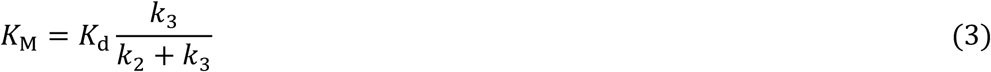

**Figure 3.**
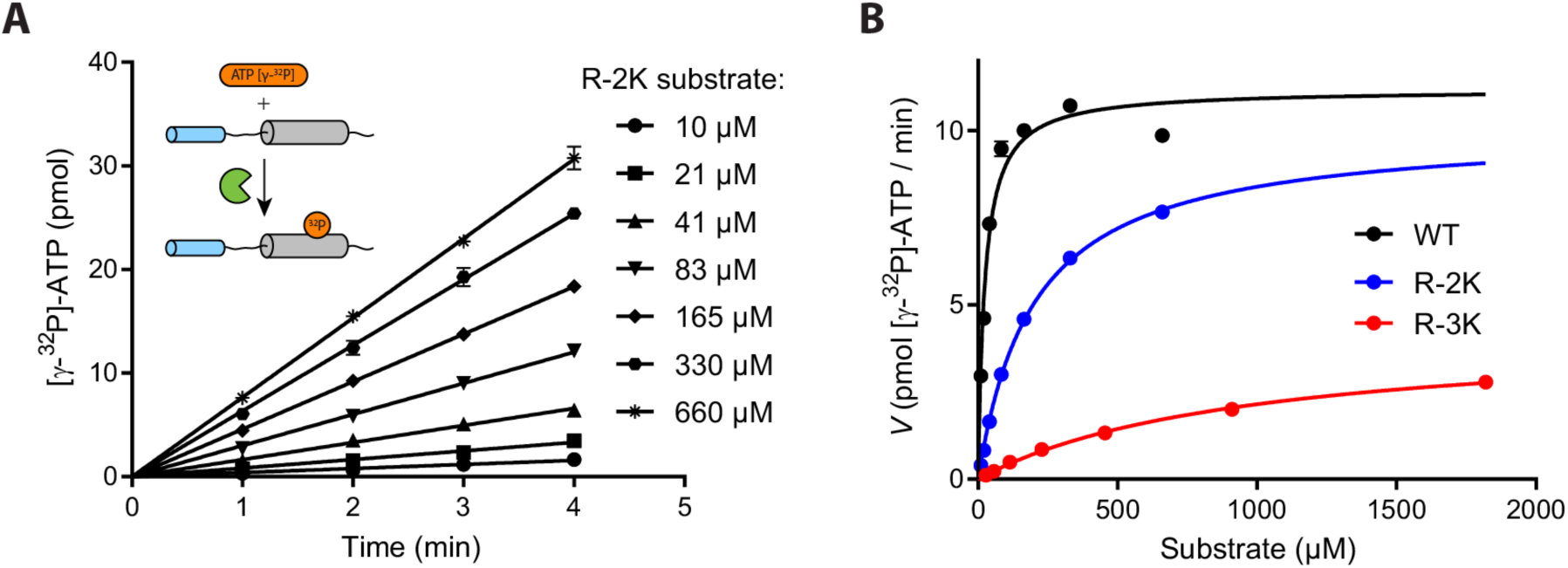
Quantifying the quality of PKA substrate variants. (A) Raw data from a steady-state kinetic experiment performed at 1 nM PKAc and an indicated concentration of the R-2K substrate. The inset shows a schematic cartoon of the experiment. (B) Steady-state kinetic data of 3 substrates fitted by a Michaelis-Menten model.

*K*_M_ is expected to be smaller than *K*_d_ when the steady-state reaction is limited by product dissociation (*k*_3_ < *k*_2_). In turn, this suggests tethered and untethered phosphorylation will have different half saturation points, as kinases are often limited by product dissociation.^25,26^

The *k*_cat_ values decreased slightly from 37.3 s^-1^ in WT to 33.5 s^-1^ and 14.1 s^-1^ in R-2K and R-3K, respectively. In contrast, there was a big difference in *k*_tet,max_ between the substrates: it was 8-fold faster than *k*_cat_ for WT (307 s^-1^), roughly the same for R-2K (40.0 s^-1^) and seemed to be slower (∼3 s^-1^) for R-3K. As *k*_tet,max_ corresponds to the rate of phosphotransfer it cannot be lower than *k*_cat_. The inconsistency for R-3K is likely due to the poorly defined plateau in the data for tethered R-3K, and thus we constrained *k*_tet,max_ to its lower limit equal to *k*_cat_ upon fitting of the R-3K data. To test how reactions with widely different rates of phosphotransfer can converge on similar steady-state turnover rates, we simulated *k*_cat_ as a function of *k*_tet,max_ for different values of the product dissociation rate constant (Fig. 4A) as below:

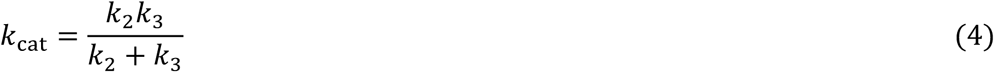

**Figure 4.**
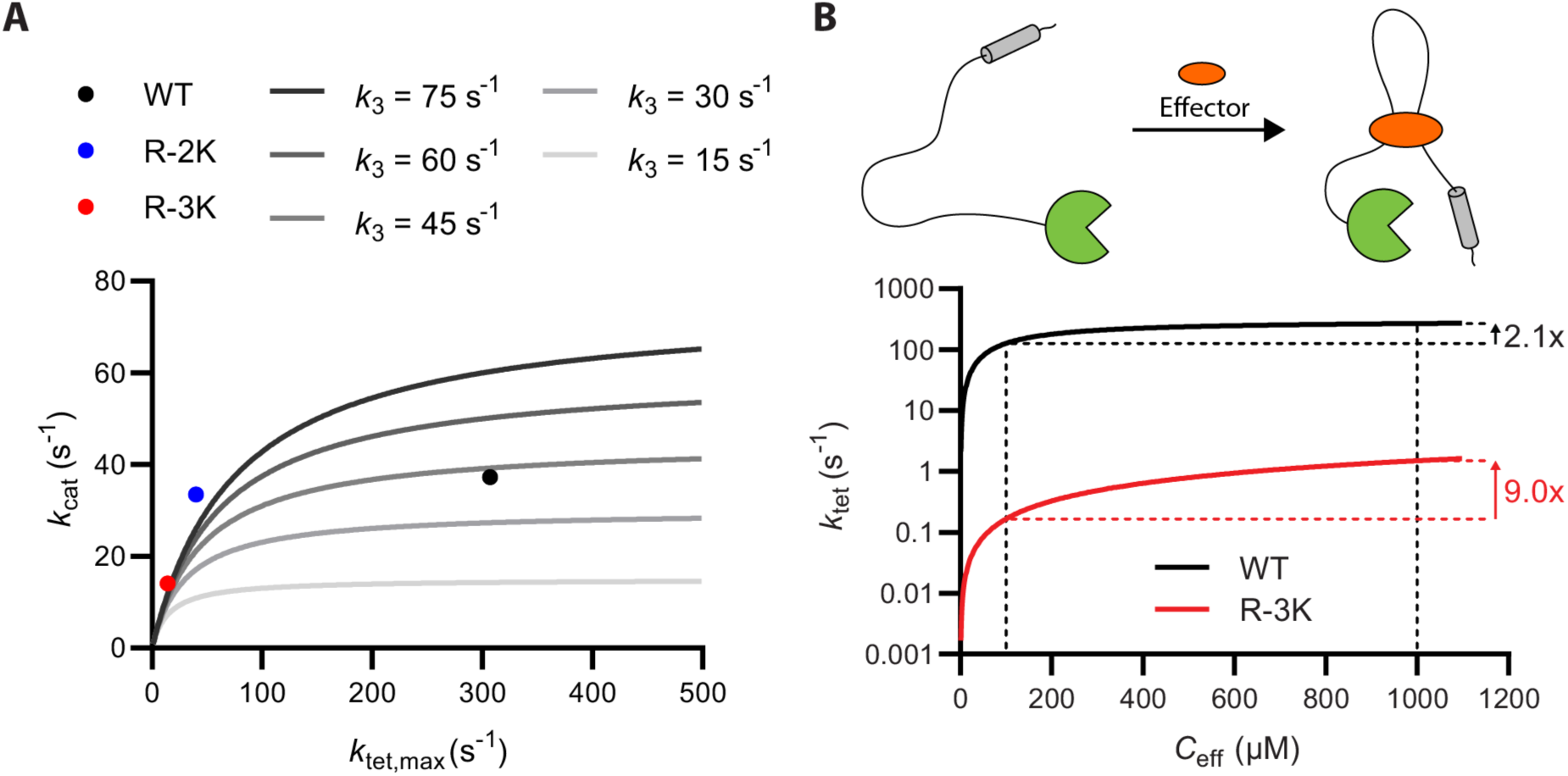
Simulated predictions for tethered catalysis. (A) Simulations of *k*_cat_ from *k*_tet,max_ and *k*_3_ based on Equation 4 in a product release limited reaction. At high k_tet,max_ values, k_cat_ is decoupled from a the rate of the tethered reaction and may disguise large differences in the tethered phosphorylation rate. (B) Fitting curves from Figure 2C (WT) and 2E (R-3K) show unequal increase in *k*_tet_ (2.1x and 9.0x, respectively) upon increasing *C*_eff_ from 100 to 1000 µM.

Fig. 4A shows that *k*_cat_ is largely independent of the rate of phospho-transfer, when the steady-state reaction is limited by product dissociation. The observed values of *k*_cat_ can thus roughly be explained assuming the same rate of product dissociation. Product dissociation rates will be similar if ADP dissociation is limiting, but may differ for the peptides, so a perfect fit is not expected. Crucially, however, Fig. 4A shows that steady-state kinetics only have a limited ability to predict the rate of a tethered reaction, and by extension it cannot predict the relative substrate usage in signalling events that only require a single intra-complex phosphorylation.

## Discussion

We have developed a reductionistic model for phosphorylation by a tethered kinase that mimics key features of kinases that contain disordered linkers or bind flexible scaffolding proteins. The model system omits the complexity of natural signalling complexes, but in return allows systematic variation of the linker architecture and substrate. This allowed us to demonstrate that phosphorylation can be enhanced dramatically by tethering, and this enhancement depends strongly on the linker architecture. We have shown that single-turnover tethered phosphorylation can be described by a Michaelis-Menten like dependence on the effective concentration, but that the kinetic parameters can only partially be described from steady-state kinetics.

We have shown that the rate of tethered phosphorylation reaches a plateau at high effective concentrations, but is such saturation relevant for kinase reactions in natural signalling complexes? Saturation occurs when the effective substrate concentration exceeds the affinity of the kinase. The affinity of kinases for short substrate motifs can typically reach low μM values, although many biologically important substrates are far from the optimal sequence motif and may have even lower affinities. Model systems and computational modelling suggest that the effective concentrations in protein complexes reach the low mM range.^12,20,27^ This suggests that signalling complexes can reach the saturation regime, where e.g. the shortening of a linker will no longer enhance the reaction.

Saturation at high effective concentrations provides a mechanism for how changes in the linker architecture can change the relative rate of tethered substrates, and thus alter the substrate usage of a kinase. For example, consider two adjacent phosphorylation sites corresponding to the WT and R-3K substrates tethered at the same effective concentration (Fig. 4B). If the scaffolding protein changes structure due to e.g. effector binding or alternative splicing, the effective concentration may change from 100 μM to 1 mM, corresponding to GS-linkers of 257 and 53 residues, respectively. These values roughly match the difference in linkers between the α and β−isoforms of CaMKII.^28^ As the WT substrate is near saturation, the phosphorylation rate of R-3K will be enhanced 4.3-fold more. This suggests that changes in linker architecture that increase effective concentrations shift relative substrate usage towards low affinity substrates.

Tethered kinases are often part of switch-like functions, where the biological effect only requires phosphorylation of few substrates. Examples include autophosphorylation of kinases such as CaMKII or activation by transphosphorylation of receptor tyrosine kinases. Furthermore, activatory stimuli are often transient such as e.g. the brief burst of Ca^2+^ triggered by opening Ca^2+^-channels. In such switches, the number of substrates processed per time is less important than the time required for the first phosphorylation. Yet steady-state kinetics remains the main mode of evaluating substrates, even if the native physiological context is a tethered complex. We show that steady-state parameters can disguise big differences in the rate of tethered phosphorylation: The WT and R-2K substrates had a 8-fold difference in *k*_tet,max_ even though the *k*_cat_ values are similar. This occurs because product dissociation limits the steady-state reaction, but not the phosphorylation of tethered substrate. The latter is more relevant to tethered kinases with switch-like functions, which suggests that substrate properties should be evaluated in single turnover experiments.

Kinases are often tethered to substrates via non-covalent interactions, which suggests that dissociation and rebinding should be considered. The model proposed here only considers a static connection, even though the fitted data uses a non-covalent connection between kinase and substrate. We suggest that the interaction can be approximated as static, when its lifetime is much longer than the catalytic cycle. As such our model system describes high-affinity scaffolding interactions in addition to linkers that are truly static, such as in kinase auto-phosphorylation reactions. For weaker interactions, single-turnover and steady-state kinetic models start to blend and will accordingly be more complex. We believe that static model will provide a baseline for such efforts.

During the preparation of this manuscript, a pre-print appeared with similar aims, but very different results.^29^ In agreement with our results, Speltz and Zalatan found that for tethered PKAc shorter linkers increased the rate of phosphorylation. Despite the similarity between the model systems used, the reaction occurred in a different kinetic regime with a maximal *k*_tet_ of ∼0.002 s^-1^. The unexpectedly slow phosphorylation rate was explained by effective concentrations of ∼80 nM, which is 50.000-fold below prediction from a simple geometric model.^29^ The systems differ mainly in the substrate and the linkers used. It is possible, that the kinetics simply reflect the different substrates, as subtle differences in the motif may have a big effect on the rate of the tethered reaction. Additionally, Speltz and Zalatan use short linkers that also contain a folded domain. Likely, the discrepancy between experimental and predicted values shows the limitation of simple polymer models for short linkers. Such systems can easily become sterically limited by the excluded volume from folded domains and sequence-specific interactions that lead to internal friction. In contrast, long disordered linkers allow almost free orientation of domains,^30^ and are likely to provide the most robust architecture for engineered scaffolding proteins.

Tethered reactions are not just important for kinases. Tethering is arguably even more important for phosphatases as the substrate motif contributes little specificity. Instead, phosphatases are regulated by anchoring proteins, and e.g. protein phosphatase-1 has at least 200 different regulatory subunits that target it to different substrates.^31^ These anchoring interactions likely display a similar dependence for linker architecture as seen here. Tethering is also increasingly used as a therapeutic intervention strategy. Enzymes tethering is important in e.g. PROTACs that target proteins for degradation by tethering them to ubiquitin ligase.^32^ Tethering has also been used pharmacologically to enhance other classes of enzymes including e.g. proteases^33^ and phosphatases.^34^ Such connector molecules will likely display a similar dependence on connection to the systems studied here, and mechanistic studies of tethered catalysis is thus both of fundamental importance to understand signalling pathways and to optimize pharmaceutical intervention strategies.

## Acknowledgments

This work was supported by grants to M.K. from the “Young Investigator Program” of the Villum Foundation, the AIAS COFUND program funded by the EU FP7 programme (Agreement no. 754513) and PROMEMO – Center for Proteins in Memory, a Center of Excellence funded by the Danish National Research Foundation (grant number DNRF133). We would like to thank Xavier Warnet and Sujata Mahapatra for critical comments to the manuscript.

