## Supplemental Information for "Intrinsically disordered linkers control tethered kinases via effective concentration"

### Protein sequences

**Color coding:**

- 6xHis-tag
- Thrombin cleavage sequence, // marks the cleavage site
- MBD2 dimerization domain
- p66α dimerization domain
- GCTAGC (AS) - NheI restriction site
- $(GS)_n$ – variable-length GS linker; n = 1, 10, 30 or 60
- GGTACC (GT) - KpnI restriction site
- PKA substrate motif, catalytic Ser shown in **bold**

#### 1. PKAc

MGSSHHHHHHSSGLVPR//GSHMGNAAAAKKGSEQESVKEFLAKAKEDFLKKWETPSQNTAQLDQFDRIKTLGT
GSFGRVMLVKHKESGNHYAMKILDKQKVVKLKQIEHTLNEKRILQAVNFPFLVKLEFSFKDNSNLYMVMEYVAGG
EMFSHLRRIGRFSEPHARFYAAQIVLTFEYLHSLDLIYRDLKPENLLIDQQGYIQVTDFGFAKRVKGRTWTLCGTPE
YLAPEIILSKGYNKAVDWWALGVLIYEMAAGYPPFFADQPIQIYEKIVSGKVRFPSHFSSDLKDLLRNLLQVDLTKRF
GNLKNGVNDIKNHKWFATTDWIAIYQRKVEAPFIPKFKGPGDTSNFDDYEEEEIRVSINEKCGKEFTEF

#### 2. MBD2-$(GS)_n$-PKAc

MGSSHHHHHHSSGLVPR//GSHMVTDEDIRKQEERAQQVRKKLEEALMADAS$(GS)_n$GTGNAAAAKKGSEQESVKE
FLAKAKEDFLKKWETPSQNTAQLDQFDRIKTLGTGSFGRVMLVKHKESGNHYAMKILDKQKVVKLKQIEHTLNEK
RILQAVNFPFLVKLEFSFKDNSNLYMVMEYVAGGEMFSHLRRIGRFSEPHARFYAAQIVLTFEYLHSLDLIYRDLKP
ENLLIDQQGYIQVTDFGFAKRVKGRTWTLCGTPEYLAPEIILSKGYNKAVDWWALGVLIYEMAAGYPPFFADQPIQI
YEKIVSGKVRFPSHFSSDLKDLLRNLLQVDLTKRFGNLKNGVNDIKNHKWFATTDWIAIYQRKVEAPFIPKFKGPG
DTSNFDDYEEEEIRVSINEKCGKEFTEF

#### 3. p66α -$(GS)_n$-WT substrate

MGSSHHHHHHSSGLVPR//GSHMTSPEERERMIKQLKEELRLEEAKLVLLKKLRQSQIQKEATAQKAS$(GS)_n$GTPG
SGSGSGSLRRA**S**LGGGGGY

#### 4. p66α -$(GS)_n$-R-2K substrate

MGSSHHHHHHSSGLVPR//GSHMTSPEERERMIKQLKEELRLEEAKLVLLKKLRQSQIQKEATAQKAS$(GS)_n$GTPG
SGSGSGSLRKA**S**LGGGGGY

#### 5. p66α -$(GS)_n$-R-3K substrate

MGSSHHHHHHSSGLVPR//GSHMTSPEERERMIKQLKEELRLEEAKLVLLKKLRQSQIQKEATAQKAS$(GS)_n$GTPG
SGSGSGSLKRA**S**LGGGGGY

### Supplementary figures

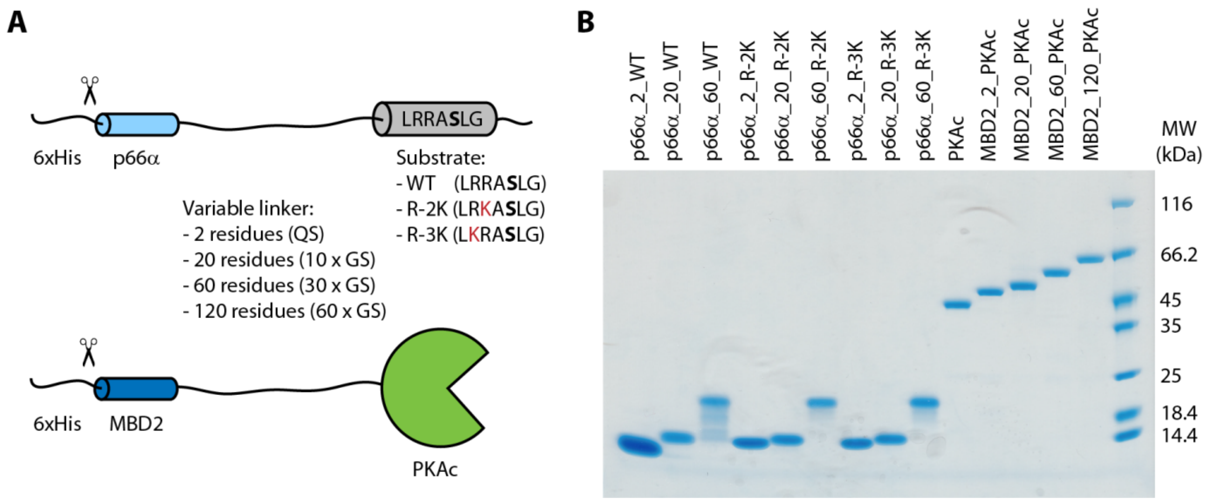


**Fig. S1: Protein variants used in this study.** (A) Schematic representation of the protein variants used to form the tethered kinase substrate complex. Both constructs contain variable GS-linker. (B) SDS-PAGE gel of purified proteins. The substrate with a 120 residue GS linker could not be purified to sufficient purity.

Next four pages:

**Fig. S2: Raw data from quench-flow experiments.**

### PKA & S WT

**Linker length:**

#### 1 mM ATP+ 5 µM PKA & S

#### 0.1 mM ATP+ 0.5 µM PKA & S

**20**

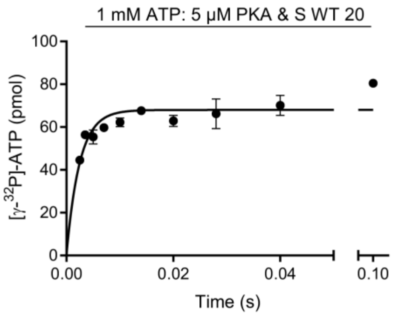


| Best-fit values | |
|---|---|
| Y0 | = 0.000 |
| Plateau | 68.00 |
| K | 406.6 |
| Tau | 0.002460 |
| Half-time | 0.001705 |
| Span | = 68.00 |
| Std. Error | |
| Plateau | 1.718 |
| K | 50.41 |

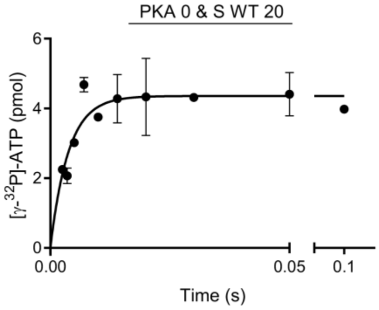


| | |
|---|---|
| Y0 | = 0.000 |
| Plateau | 4.355 |
| K | 263.9 |
| Tau | 0.003790 |
| Half-time | 0.002627 |
| Span | = 4.355 |
| Std. Error | |
| Plateau | 0.1957 |
| K | 47.04 |

**40**

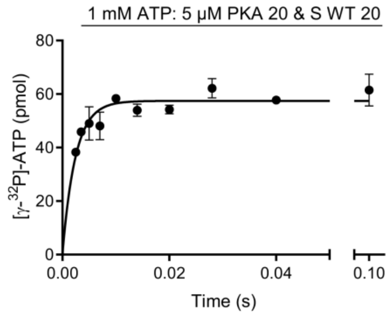


| Best-fit values | |
|---|---|
| Y0 | = 0.000 |
| Plateau | 57.44 |
| K | 411.2 |
| Tau | 0.002432 |
| Half-time | 0.001686 |
| Span | = 57.44 |
| Std. Error | |
| Plateau | 1.356 |
| K | 46.58 |

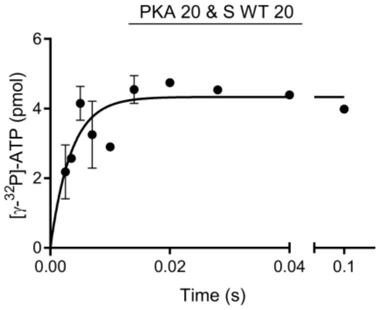


| | |
|---|---|
| Y0 | = 0.000 |
| Plateau | 4.331 |
| K | 287.6 |
| Tau | 0.003477 |
| Half-time | 0.002410 |
| Span | = 4.331 |
| Std. Error | |
| Plateau | 0.2459 |
| K | 58.18 |

**60**

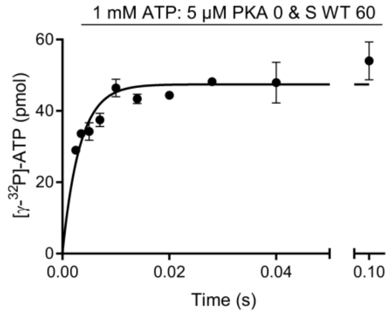


| Best-fit values | |
|---|---|
| Y0 | = 0.000 |
| Plateau | 47.44 |
| K | 306.7 |
| Tau | 0.003260 |
| Half-time | 0.002260 |
| Span | = 47.44 |
| Std. Error | |
| Plateau | 1.275 |
| K | 34.45 |

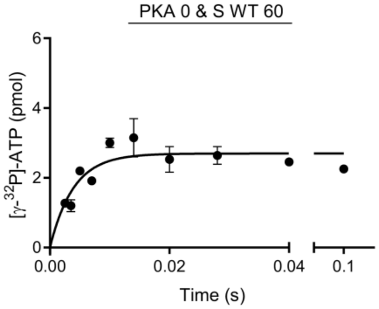


| | |
|---|---|
| Y0 | = 0.000 |
| Plateau | 2.694 |
| K | 254.7 |
| Tau | 0.003927 |
| Half-time | 0.002722 |
| Span | = 2.694 |
| Std. Error | |
| Plateau | 0.1361 |
| K | 47.63 |

**80**

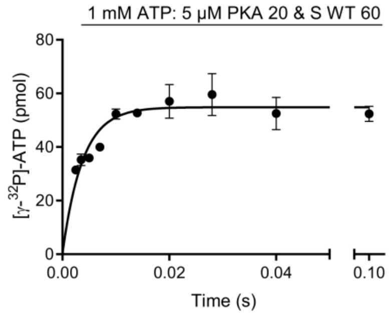


| Best-fit values | |
|---|---|
| Y0 | = 0.000 |
| Plateau | 54.85 |
| K | 267.7 |
| Tau | 0.003735 |
| Half-time | 0.002589 |
| Span | = 54.85 |
| Std. Error | |
| Plateau | 1.508 |
| K | 29.40 |

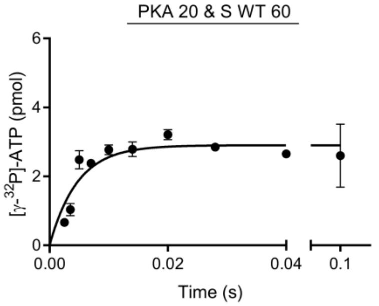


| Best-fit values | |
|---|---|
| Y0 | = 0.000 |
| Plateau | 2.900 |
| K | 212.2 |
| Tau | 0.004713 |
| Half-time | 0.003267 |
| Span | = 2.900 |
| Std. Error | |
| Plateau | 0.1629 |
| K | 40.88 |

**120**

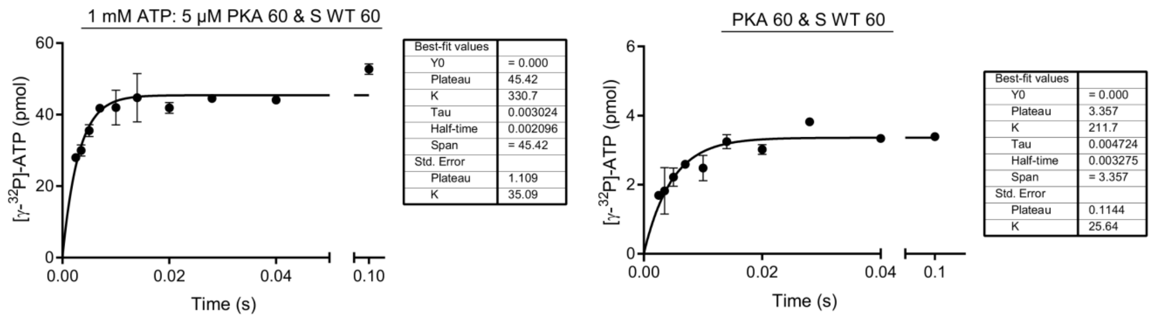


**180**

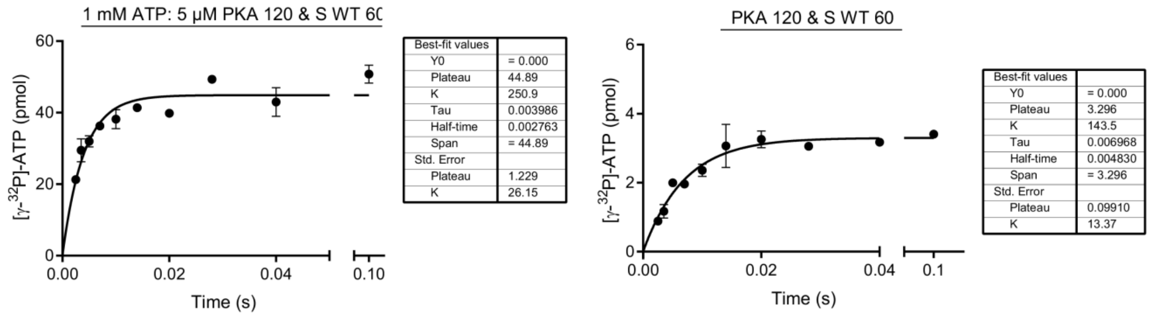


**not linked**

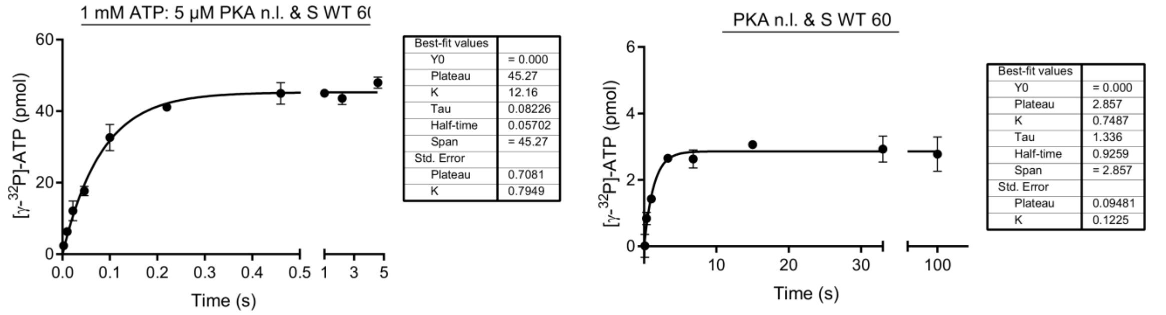

| | PKA & S R-2K | PKA & S R-3K |
|---|---|---|
| **Linker length:** | **0.1 mM ATP+ 0.5 µM PKA & S** | **0.1 mM ATP+ 0.5 µM PKA & S** |

**20**

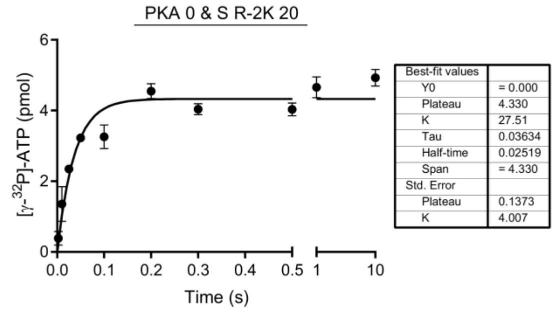


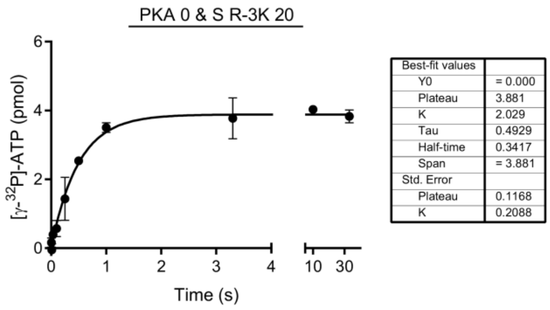


**40**

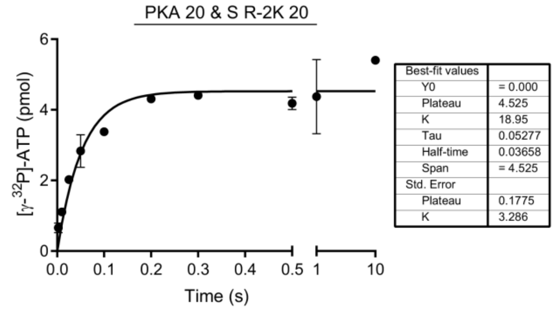


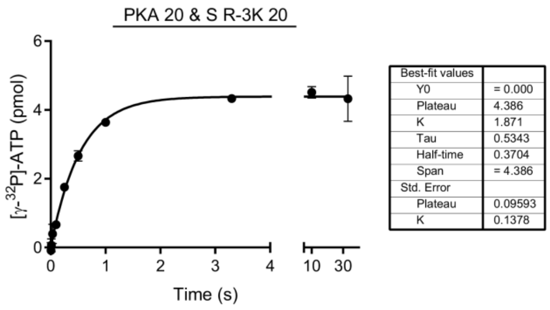


**60**

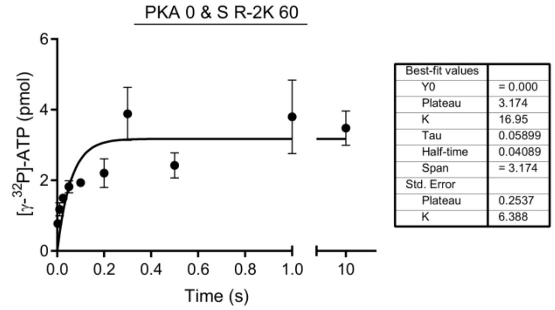


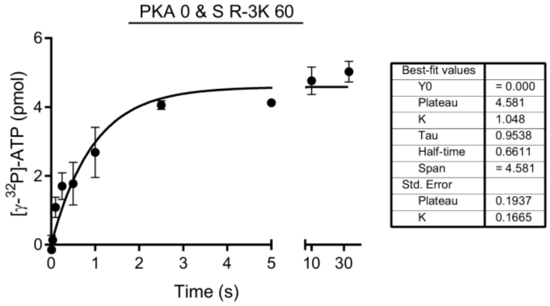


**80**

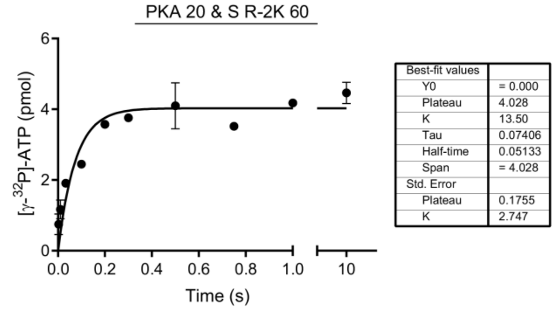


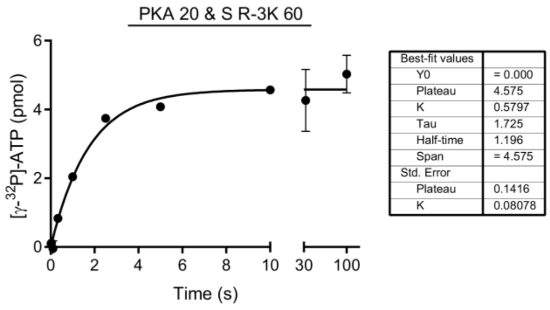

**120**

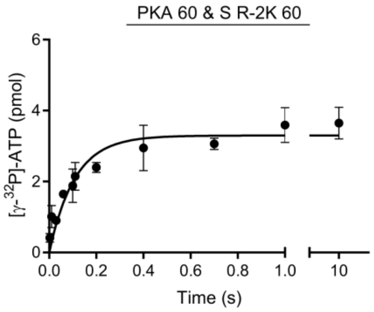


| Best-fit values | |
|---|---|
| Y0 | = 0.000 |
| Plateau | 3.297 |
| K | 9.214 |
| Tau | 0.1085 |
| Half-time | 0.07522 |
| Span | = 3.297 |
| Std. Error | |
| Plateau | 0.1629 |
| K | 1.538 |

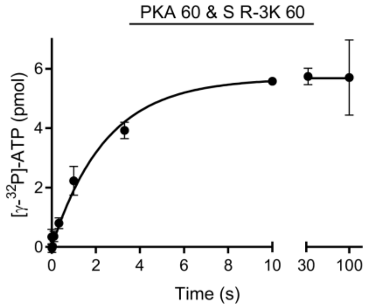


| Best-fit values | |
|---|---|
| Y0 | = 0.000 |
| Plateau | 5.680 |
| K | 0.4094 |
| Tau | 2.442 |
| Half-time | 1.693 |
| Span | = 5.680 |
| Std. Error | |
| Plateau | 0.1846 |
| K | 0.05113 |

**180**

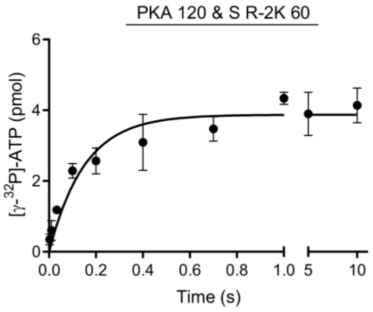


| Best-fit values | |
|---|---|
| Y0 | = 0.000 |
| Plateau | 3.872 |
| K | 6.625 |
| Tau | 0.1510 |
| Half-time | 0.1046 |
| Span | = 3.872 |
| Std. Error | |
| Plateau | 0.1713 |
| K | 1.276 |

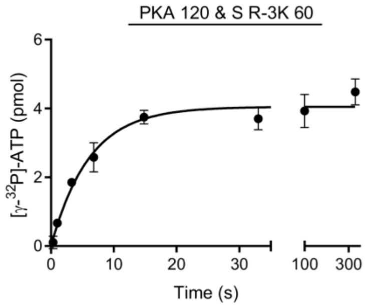


| Best-fit values | |
|---|---|
| Y0 | = 0.000 |
| Plateau | 4.046 |
| K | 0.1644 |
| Tau | 6.081 |
| Half-time | 4.215 |
| Span | = 4.046 |
| Std. Error | |
| Plateau | 0.1284 |
| K | 0.02013 |

**not linked**

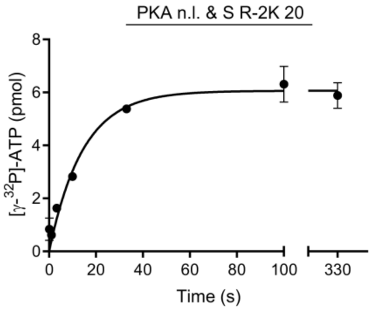


| Best-fit values | |
|---|---|
| Y0 | = 0.000 |
| Plateau | 6.061 |
| K | 0.06950 |
| Tau | 14.39 |
| Half-time | 9.973 |
| Span | = 6.061 |
| Std. Error | |
| Plateau | 0.2303 |
| K | 0.01012 |

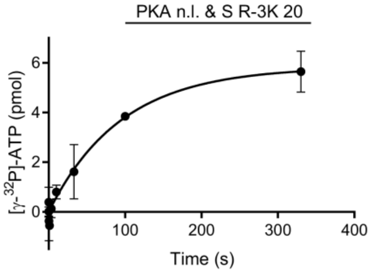


| Best-fit values | |
|---|---|
| Y0 | = 0.000 |
| Plateau | 5.833 |
| K | 0.01056 |
| Tau | 94.70 |
| Half-time | 65.64 |
| Span | = 5.833 |
| Std. Error | |
| Plateau | 0.4792 |
| K | 0.002252 |

### PKA & S R-3K

#### 1 mM ATP+ 5 µM PKA & S

**60**

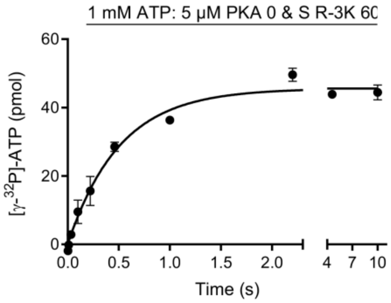


| Best-fit values | |
|---|---|
| Y0 | = 0.000 |
| Plateau | 45.58 |
| K | 1.972 |
| Tau | 0.5071 |
| Half-time | 0.3515 |
| Span | = 45.58 |
| Std. Error | |
| Plateau | 1.113 |
| K | 0.1746 |

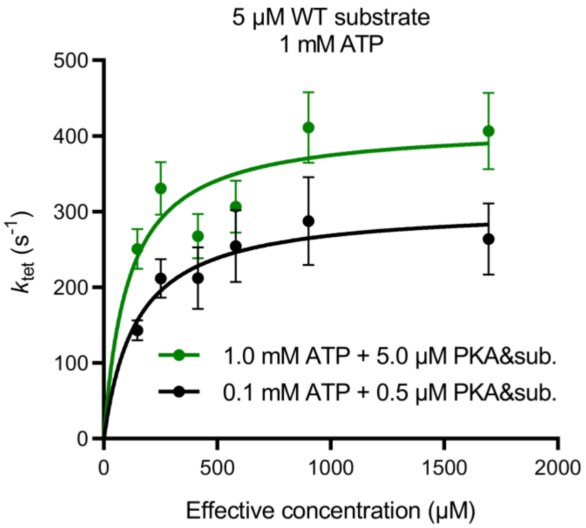


**Fig. S3: ATP dependence of the tethered reaction.** The ATP dependence of the tethered phosphorylation was tested at 1 mM ATP, whereas all other experiments were conducted at 100 µM. The amount of [γ-$^{32}$P]ATP was already at maximally permitted level, so the protein concentration was also increased 10-fold to preserve the same signal to noise.

### Derivations of rate equations

#### *Tethered system*

We consider a catalytical model where product release limits steady-state reaction rates and where phosphorylation and product release are two irreversible steps.

Moreover, we use saturating ATP concentrations and assume $k_{\text{ATP binding}} >> k_{\text{cat}}$. Thus, we define the following states:

- O = open tethered system, bound ATP
- C = closed tethered system, bound ATP
- CP = closed, phosphorylated tethered system, bound ADP
- OP = open tethered system

The tethered system is composed of two interacting partners, and closure of this system is governed by effective concentration, $C_{\text{eff}}$:

$$O \underset{k_{-1}}{\overset{k_1 C_{eff}}{\longleftrightarrow}} C \overset{k_2}{\rightarrow} CP \overset{k_3}{\rightarrow} OP$$

The Law of Mass Action applied to the model leads to the following system of nonlinear reaction equations:

$$\frac{d[O]}{dt} = -k_1 C_{eff}[O] + k_{-1}[C]$$

$$\frac{d[C]}{dt} = k_1 C_{eff}[O] - (k_{-1} + k_2)[C]$$

$$\frac{d[CP]}{dt} = k_2[C] - k_3[CP]$$

$$\frac{d[OP]}{dt} = k_3[CP]$$

In single turnover experiments both closed and open phosphorylated products are measured, hence:

$$P = CP + OP$$

$$\frac{d[P]}{dt} = \frac{d[CP]}{dt} + \frac{d[OP]}{dt} = k_2[C]$$

From the conservation law, total concentration of the tethered system is constant:

$$\frac{d[O]}{dt} + \frac{d[C]}{dt} + \frac{d[CP]}{dt} + \frac{d[OP]}{dt} = 0$$

$$[O] + [C] + [CP] + [OP] = [E]_\mathrm{T}$$

$$[O] = [E]_\mathrm{T} - [C] - [P]$$

Rapid equilibrium assumption for the open/closed complex:

$$\frac{d[O]}{dt} = 0$$

$$k_1 C_{eff}[O] = k_{-1}[C]$$

$$k_1 C_{eff}([E]_\mathrm{T} - [C] - [P]) = k_{-1}[C]$$

$$k_{-1}[C] + k_1 C_{eff}[C] = k_1 C_{eff}[E]_\mathrm{T} - k_1 C_{eff}[P]$$

$$[C] = \frac{k_1 C_{eff}[E]_\mathrm{T} - k_1 C_{eff}[P]}{k_{-1} + k_1 C_{eff}} = \frac{C_{eff}[E]_\mathrm{T} - C_{eff}[P]}{\frac{k_{-1}}{k_1} + C_{eff}}$$

Given $K_\mathrm{d} = \frac{k_{-1}}{k_1}$:

$$[C] = \frac{C_{eff}[E]_\mathrm{T} - C_{eff}[P]}{K_d + C_{eff}}$$

Substituting $[C]$ into the product formation equation:

$$\frac{d[P]}{dt} = k_2[C] = k_2 \frac{C_{eff}[E]_\mathrm{T} - C_{eff}[P]}{K_d + C_{eff}}$$

Integrate product formation rate:

$$\frac{d[P]}{dt} = -\frac{k_2 C_{eff}}{K_d + C_{eff}}[P] + \frac{k_2 C_{eff}[E]_\mathrm{T}}{K_d + C_{eff}}$$

$$\frac{1}{-\frac{k_2 C_{eff}}{K_d + C_{eff}}[P] + \frac{k_2 C_{eff}[E]_\mathrm{T}}{K_d + C_{eff}}} d[P] = dt$$

$$\int_0^{[P]} \frac{1}{-\frac{k_2 C_{eff}}{K_d + C_{eff}}[P] + \frac{k_2 C_{eff}[E]_\mathrm{T}}{K_d + C_{eff}}} d[P] = \int_0^t dt$$

$$-\frac{K_d + C_{eff}}{k_2 C_{eff}} \ln\left|-\frac{k_2 C_{eff}}{K_d + C_{eff}}[P] + \frac{k_2 C_{eff}[E]_\mathrm{T}}{K_d + C_{eff}}\right| + \frac{K_d + C_{eff}}{k_2 C_{eff}} \ln\left|-\frac{k_2 C_{eff}}{K_d + C_{eff}} \cdot 0 + \frac{k_2 C_{eff}[E]_\mathrm{T}}{K_d + C_{eff}}\right| = t - 0$$

$$\ln\left|\frac{k_2 C_{eff}([E]_\mathrm{T} - [P])}{K_d + C_{eff}}\right| - \ln\left|\frac{k_2 C_{eff}[E]_\mathrm{T}}{K_d + C_{eff}}\right| = -\frac{k_2 C_{eff}}{K_d + C_{eff}}t$$

$$\ln\left|\frac{[E]_\mathrm{T} - [P]}{[E]_\mathrm{T}}\right| = -\frac{k_2 C_{eff}}{K_d + C_{eff}}t$$

$$\frac{[E]_\mathrm{T} - [P]}{[E]_\mathrm{T}} = e^{-\frac{k_2 C_{eff}}{K_d + C_{eff}}t}$$

Formation of phosphorylated product is described by the following equation:

$$[P] = [E]_\mathrm{T}\left(1 - e^{-\frac{k_2 C_{eff}}{K_d + C_{eff}}t}\right)$$

Phosphorylation rates in the tethered system obtained from quench flow measurements ($k_{\mathrm{tet}}$) are dependent on effective concentration:

$$k_{tet} = \frac{k_2 C_{eff}}{K_d + C_{eff}}$$

***Untethered system***

We consider a catalytical model where product release limits steady-state reaction rates and where phosphorylation and product release are two irreversible steps.

$$S + E \underset{k_{-1}}{\overset{k_1}{\longleftrightarrow}} ES \overset{k_2}{\rightarrow} EP \overset{k_3}{\rightarrow} E + P$$

The Law of Mass Action applied to the model leads to the following system of nonlinear reaction equations:

$$\frac{d[S]}{dt} = -k_1[S][E] + k_{-1}[ES]$$

$$\frac{d[E]}{dt} = -k_1[S][E] + k_{-1}[ES] + k_3[EP]$$

$$\frac{d[ES]}{dt} = k_1[S][E] - (k_{-1} + k_2)[ES]$$

$$\frac{d[EP]}{dt} = k_2[ES] - k_3[EP]$$

$$\frac{d[P]}{dt} = k_3[EP]$$

From the conservation law for the enzyme, total enzyme concentration is constant:

$$\frac{d[E]}{dt} + \frac{d[ES]}{dt} + \frac{d[EP]}{dt} = 0$$

$$[E] + [ES] + [EP] = [E]_0$$

$$[E] = [E]_0 - [ES] - [EP]$$

Rapid equilibrium assumption:

$$\frac{d[S]}{dt} = 0$$

$$k_1[S][E] = k_{-1}[ES]$$

$$k_1[S]([E]_0 - [ES] - [EP]) = k_{-1}[ES]$$

$$k_1[S][E]_0 - k_1[S][EP] = k_{-1}[ES] + k_1[S][ES]$$

$$[ES] = \frac{k_1[S][E]_0 - k_1[S][EP]}{k_1[S] + k_{-1}}$$

Quasi-steady-state approximation of the $[EP]$ complex:

$$\frac{d[EP]}{dt} = 0$$

$$k_2[ES] - k_3[EP] = 0$$

$$\frac{k_3[EP]}{k_2} = [ES]$$

$$\frac{k_3[EP]}{k_2} = \frac{k_1[S][E]_0 - k_1[S][EP]}{k_1[S] + k_{-1}}$$

$$k_1k_3[S][EP] + k_{-1}k_3[EP] = k_1k_2[S][E]_0 - k_1k_2[S][EP]$$

$$[EP]([S]k_1(k_2 + k_3) + k_{-1}k_3) = k_1k_2[S][E]_0$$

$$[EP] = \frac{k_1k_2[S][E]_0}{[S]k_1(k_2 + k_3) + k_{-1}k_3}$$

$$[EP] = \frac{\dfrac{k_2[S][E]_0}{k_2 + k_3}}{[S] + \dfrac{k_{-1}k_3}{k_1(k_2 + k_3)}}$$

Given $K_\mathrm{d} = \frac{k_{-1}}{k_1}$:

$$[EP] = \frac{\dfrac{k_2[S][E]_0}{k_2 + k_3}}{[S] + K_\mathrm{d}\dfrac{k_3}{k_2 + k_3}}$$

Finally, substituting $[EP]$ into the product formation equation:

$$\frac{d[P]}{dt} = k_3[EP] = \frac{\dfrac{k_2k_3}{k_2 + k_3}[S][E]_0}{[S] + K_\mathrm{d}\dfrac{k_3}{k_2 + k_3}}$$

Hence:

$$k_{cat} = \frac{k_2k_3}{k_2 + k_3}$$

$$K_\mathrm{M} = K_\mathrm{d}\frac{k_3}{k_2 + k_3}$$
